# Single electrically induced epileptic afterdischarge triggers synaptic weakening, AMPA receptor endocytosis and shrinkage of dendritic spines in the hippocampus

**DOI:** 10.1101/2025.04.15.649044

**Authors:** Ana Paula Crestani, Leonardo Rakauskas Zacharias, Danilo Benette Marques, Flavia Faria Formagio Fonseca, Rafael Naime Ruggiero, João Pereira Leite

## Abstract

Afterdischarge (AD) is an experimental model of electrically induced seizures. When induced in limbic structures, AD is known to induce acute behavioral alterations and flattening of local field potentials that persist for a few minutes. However, impairments in more complex cognitive processes such as learning and memory, can last for hours after the seizure. Considering the mismatch between ephemeral postictal electrophysiological changes and long-lasting cognitive impairments, synaptic plasticity emerges as a possible mechanism that integrates these findings. Therefore, we used a multilevel approach to evaluate the effects of a single seizure on synaptic plasticity. First, we showed that a single AD-eliciting stimulation at the perforant pathway-dentate gyrus (PP-DG) synapse causes long-lasting decrease of evoked postsynaptic potentials. Then, we observed that this functional alteration was accompanied by shrinkage of mushroom-shaped spines in the DG. Lastly, reduced levels of p(Ser845)-GluA1 and GluA1 subunits of AMPA receptors (AMPARs) were found in the hippocampal postsynaptic density. Together, our data suggest that a seizure induces a long-term depression (LTD)-like synaptic plasticity phenomenon, which is characterized by synaptic weakening, AMPAR endocytosis, and dendritic spines shrinkage. Furthermore, the postictal changes in synaptic plasticity observed here align with the course of cognitive impairments often reported in other studies.

## 1. INTRODUCTION

Postictal states are typically characterized by mental confusion, impaired learning ability, and a reduced likelihood of immediate subsequent seizures. These transient neurological states often follow acute seizures and can last from minutes to hours, despite the resolution of the seizure itself (1).

Afterdischarge (AD) is a well-established model of electrically induced acute seizures that can be triggered in specific brain regions. In freely moving animals, AD elicits focal impaired awareness seizures when applied to limbic structures. Hippocampal AD, in particular, causes several acute behavioral alterations, such as freezing during stimulation, wet dog shake at the end of the stimulation, followed by stereotypies and hyperlocomotion shortly thereafter. Electrophysiologically, AD is characterized by an abnormal, excessive, hypersynchronous discharge of a population of neurons that persists for seconds to minutes after the cessation of electrical stimulation. This activity is typically followed by a postictal period marked by local field potential (LFP) flattening, which may last several minutes (2,3).

Although both LFP flattening and acute behavioral alterations resolve quickly after AD, impairments in more complex cognitive tasks involving learning and memory may persist for hours (2,3). This temporal dissociation suggests that LFP flattening cannot fully account to the long-lasting cognitive deficits observed after seizures. Thus, changes in synaptic plasticity may serve as a more plausible mechanism linking the ephemeral postictal electrophysiological changes to the long-lasting cognitive impairments.

While a few studies have reported transient synaptic weakening in the hippocampus following seizure induction (4,5), most have focused exclusively on electrophysiological changes, overlooking structural and molecular alterations. Also, the recurrent seizure episodes characteristic of epileptic conditions makes it difficult to isolate the brain changes triggered by the very first seizure episode, whose level of severity may be a predictor of subsequent seizures and the establishment of an epileptic condition with a worse prognosis (6,7). To our knowledge, no previous studies have thoroughly investigated the effects of a single seizure on hippocampal synaptic plasticity in freely behaving rodents.

In the present study, we aimed to address this gap by examining the effects of a single electrically induced AD on synaptic plasticity at multiple levels—functional, molecular, and structural—following an acute seizure. We found that hippocampal AD induces a lasting (>1h) synaptic weakening at perforant pathway-dentate gyrus (PP-DG) synapse, which is accompanied by a reduction in the levels of p(Ser845)-GluA1 and GluA1 subunits of AMPA receptors (AMPAR) in the postsynaptic density and a shrinkage of mushroom spines in the upper blade of the DG. These functional, structural and molecular alterations resemble long-term depression (LTD) and may underlie the cognitive impairments observed in the postictal period.

## 2. METHODS

See on-line Supplementary Information for full description of methods and statistical analysis.

### 2.1 Animals

Two-months-old male Wistar Hannover rats were used in the experiments. All procedures followed the National Council for the Control of Animal Experimentation guidelines and were approved by the Committee on Ethics in the Use of Animals (Ribeirao Preto Medical School, University of Sao Paulo, protocol: 156/2018).

### 2.2 Electrophysiological stimulation and recordings

Stereotaxic surgery was performed under anesthesia to implant a recording electrode in the DG and an ipsilateral stimulating electrode in the PP. After a week, evoked field post-synaptic potentials (fPSP) and concomitant hippocampal LFP were recorded in freely moving animals. The recording session started with a 10-min baseline (paired-pulse every 20 seconds). Next, a 10-second 50 Hz biphasic electrical stimulation was applied to elicit an AD. A successful AD induction was defined as a rhythmic, hypersynchronous discharge lasting for more than 5 seconds. After that, evoked potentials were recorded for an additional hour. Animals were then immediately euthanized to perform structural and molecular analysis. Custom scripts in MATLAB were used for electrophysiological signal processing. Sham animals were implanted but not stimulated.

### 2.3 Golgi-Cox staining and dendritic spines quantification

After performing Golgi-Cox staining, images of the secondary or tertiary branches of granule cells dendrites located in the upper blade of the DG from both dorsal and ventral hippocampus were captured with an optical microscope. Dendrite segments (∼10um) with clear spines were used for quantification (100× magnification) and classification into different spine subtypes (long thin, thin, stubby or mushroom-shaped) using Fiji software.

### 2.4 AMPAR subunits isolation and quantification

Hippocampal tissue was subcellularly fractionated to isolate postsynaptic density samples. Western blots were performed, and proteins were transferred to membranes that were incubated with antibodies for p(Ser845)-GluA1 or GluA1 receptor subunits. β-tubulin antibody was used as loading control. Biotinylated secondary antibody and streptavidin-horseradish peroxidase conjugate were used to amplify and reveal the primary antibody signal. Immunoblotting signals were quantified using ImageLab, BioRad.

## 3. RESULTS

### 3.1 Effect of a single AD-inducing stimulation on synaptic strength at the PP-DG synapse

Changes in synaptic strength are typically associated with the ability to store information (8). Here, we investigated the effects of an electrically induced hippocampal AD on synaptic strength at the PP-DG synapse (**Fig.1A)** in freely behaving rats (**Fig.1B**, *n*=10). The fPSP2 amplitude during baseline (10-min block) was compared to the 10-min blocks following AD-inducing stimulus. One-way ANOVA showed a robust difference across blocks (*F*(_6,63)_=85.85, *p*<0.0001) and Dunnett’s multiple comparison post-hoc test showed that every block following AD differed from the baseline (*p*<0.0001) (**Fig.1C,1D)**. These results indicate that an AD-inducing stimulation causes a long-lasting reduction in fPSP amplitude.

**Figure 1.**
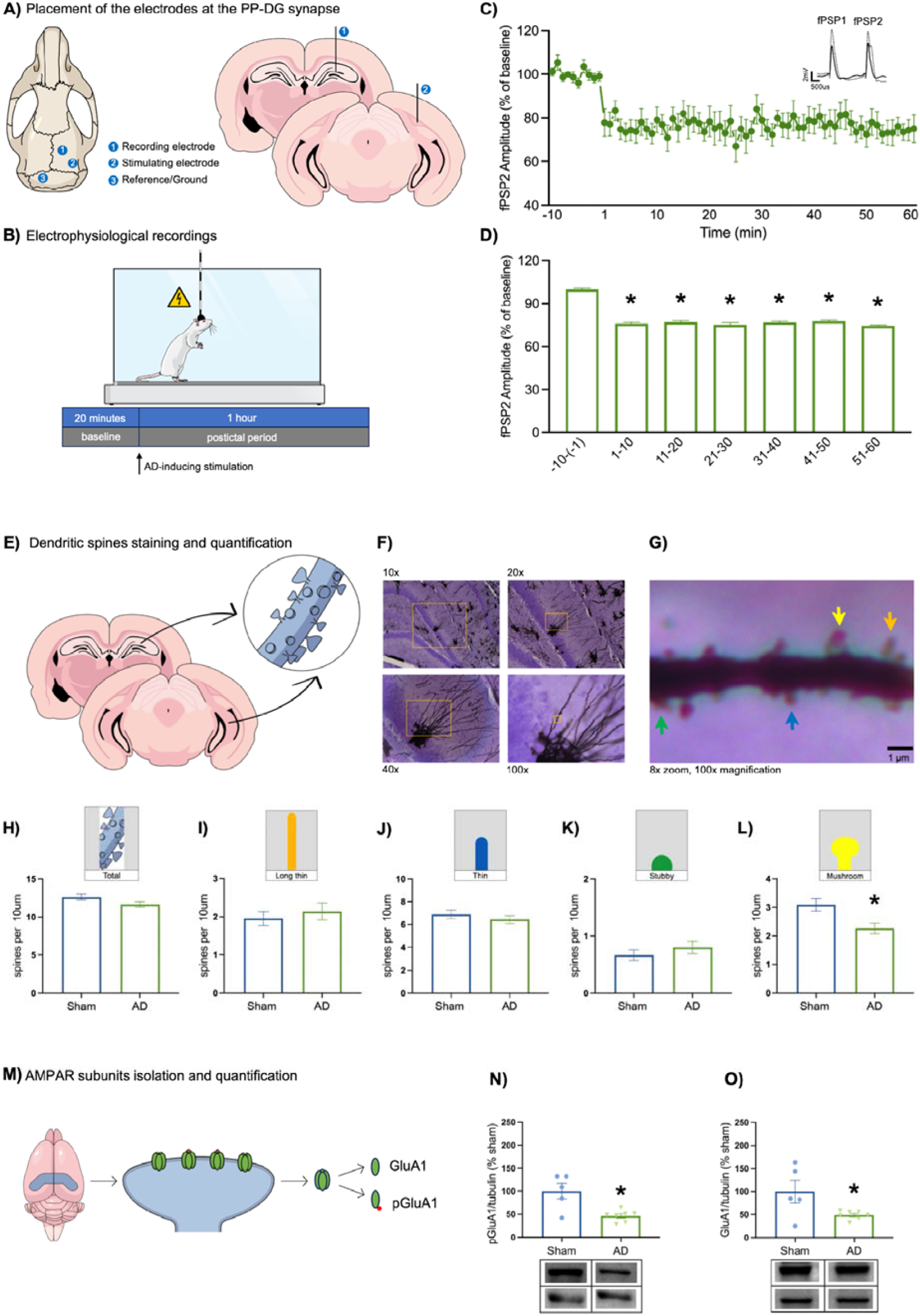
Effect of a single seizure on synaptic efficacy, dendritic spine density and AMPAR trafficking. **(A)** Schematic diagram of (1) recording electrode placement in granular cell layer of the upper blade of the DG, (2) stimulating electrode in the PP and (3) reference/ground electrode in the occipital bone. **(B)** Electrophysiological recording session timeline. Evoked potentials were recorded for a 10 min baseline. Then, AD was elicited (arrow) by applying a 10-second 50 Hz biphasic electrical stimulation. After that, evoked potentials were recorded for an additional hour. **(C)** Effects of AD-inducing stimulation on fPSP2 shown in 1-min blocks as mean ± standard error of the mean (SEM). (Inset) Representative image of the fPSP1 and fPSP2. The dashed line represents the potentials before the AD-inducing stimulation, while the solid line shows the potentials after the stimulation. **(D)** Bar graph grouping 10-min blocks as mean ± SEM. *Significant difference from the 10-min baseline block (one-way ANOVA with Dunnett’s post hoc test: *p*<0.0001). **(E)** Schematic diagram of Golgi-Cox staining and dendritic spines quantification. The density and classification of dendritic spines were performed on secondary and tertiary branches of the dendrites of granule cells located in the upper blade of the DG of both dorsal and ventral hippocampus. **(F)** Representative images of DG stained with Golgi-Cox and counterstained with Nissl. Images acquired under an optical microscope at magnification of 10× (top left), 20× (top right), 40× (bottom left), 100× (bottom right). Orange rectangles represent the field of view where the image with subsequent higher magnification was acquired. **(G)** Representative dendritic segment (10⎧m length, 8× zoom of 100× magnification image) showing spines of different morphologies. Blue arrow is pointing to a thin spine, orange arrow to a long thin spine, yellow arrow to a mushroom spine and green arrow to a stubby spine. **(H)** Total density of dendritic spines measured 1h after AD-inducing stimulation (green bar) or sham (blue bar). **(I, J, K, L)** Density of long thin **(I)**, thin **(J)**, stubby **(K)** and mushroom **(L)** spines in sham (blue bars) versus animals that underwent an AD-inducing stimulation (green bars). *Linear mixed-effects model, *p*<0.05. **(M)** Schematic diagram of AMPAR subunits isolation and quantification. Animals were euthanized 1h following AD-inducing stimulation and sham/no-stimulation animals were used as controls. **(N, O)** Expression of pGluA1- **(M)** and GluA1- **(N)** AMPARs subunits in the postsynaptic density of the hippocampus 1h after AD-inducing stimulation (green) or after no stimulation (blue). *Independent *t-*test, *p*<0.05.

### 3.2 Effect of a single AD-inducing stimulation on dendritic spine density and morphology

The structure of dendritic spines (density, size, and shape) can regulate the strength with which postsynaptic neurons respond to synaptic inputs (10). Here, morphological alterations were measured 1h after AD-inducing stimulation (*n*=4) compared to sham (*n*=4, **Fig.1E**). We used a linear mixed-effects model to assess the effects of stimulation (AD-inducing versus no-stimulation) in the number and classification of dendritic spines, accounting for repeated measurements within subjects (*n*=20 dendritic segments from each animal, **Fig.1F,1G**). Animal was included as a random intercept to model between-subject variability, and stimulation was included as a fixed effect. The model revealed a significant effect of stimulation in the mushroom-shaped spines. Specifically, animals in the AD-inducing stimulation group had significantly lower level of mushroom spines compared to the control group (*β*=−0.825, *SE*=0.282, *t*_*(158)*_=−2.92, *p*=0.004). The 95% confidence interval for the treatment effect ranged from -1.38 to -0.27, indicating a robust negative effect. The intercept (*β*=3.09) represents the estimated mean response in the control group. No differences were found between groups in the total number of dendritic spines (**Fig. 1H**), nor in the number of spines classified as long thin (**Fig.1I**), thin (**Fig. 1J**) and stubby (**Fig. 1K**).

### 3.3 Effect of a single AD-inducing stimulation on AMPAR trafficking at postsynaptic density

Synaptic weakening is often accompanied by the endocytosis of AMPARs at the postsynaptic membrane (9). Here, AMPAR subunits (p(Ser845)-GluA1 and GluA1) were quantified in subfractions of hippocampal homogenate 1h after AD-inducing stimulation (*n*=5) compared to sham (*n*=7; **Fig. 1M**). A decrease in the expression of p(Ser845)-GluA1 (independent *t*-test, *t*_*(10)*_=3.540, *p*=0.0054, **Fig. 1N**) and GluA1 (independent *t*-test, *t*_*(10)*_=2.465, *p*=0.0334, **Fig. 1O**) subunits of AMPARs were observed in the postsynaptic density fraction of the hippocampal homogenate.

## 4. DISCUSSION

In this study, we dissected the changes in synaptic plasticity triggered by a single electrically induced seizure at the PP-DG synapse of freely behaving rats. We have shown that hippocampal AD-inducing stimulation causes a long-lasting synaptic weakening at the PP-DG synapse that is accompanied by shrinkage of mushroom-shaped spines in the upper blade of the DG and reduced levels of p(Ser845)-GluA1 and GluA1 subunits of AMPARs in the hippocampal postsynaptic density.

Changes in synaptic strength measured via evoked field post-synaptic potentials (fPSPs) are often used to examine synaptic plasticity phenomena such as long-term potentiation (LTP) and LTD, in which an increase in synaptic efficacy is observed in the former and decreases in the latter (11). Our results indicate that the postictal synaptic weakening in the PP-DG synapse is long lasting (>1h) and resembles an LTD phenomenon. Similar results of postictal reduction in the evoked responses in DG were observed in gerbils with inherited epilepsy, although the effects rapidly returned to baseline (∼ 2 minutes) (5). In addition, a seizure elicited in the CA3 of rat hippocampal slices depressed CA1 responses for around 15 minutes (4), a shorter interval than we have identified. These more transitory decreases in synaptic responses compared to our data are possibly due to differences in the nature of the seizure (electrically induced versus spontaneous), brain state during seizure (awake versus anesthetized), animal species (rats versus gerbils) and synapses (PP-DG versus CA3-CA1) investigated.

The LTD-like phenomenon revealed here might explain effects frequently observed after a seizure, such as inhibition of processes that depend on synaptic strengthening. Although we did not assess whether the ability to induce LTP is compromised during the postictal synaptic weakening, other results point in this direction. Barr et al., for example, showed LTP induction was inhibited during the period of postictal depression of evoked responses (4). Furthermore, given that adjustments in synaptic strength modify the ability to store information (8), mnemonic processes linked to the potentiation of the postsynaptic response may be impaired after seizures (12,13).

Alternatively, the postictal LTD-like phenomenon can be interpreted as a negative feedback mechanism that counteracts AD-induced neural hyperexcitability, characterizing a phenomenon of homeostatic plasticity (14). Homeostatic plasticity occurs over a number of different spatial scales from the synaptic to the cell population level and can act at different rates, ranging from minutes to days. It is difficult to disentangle homeostatic (e.g., synaptic downscaling) from Hebbian (e.g. LTD) plasticity, especially since both types of plasticity can occur in parallel and have overlapping underlying mechanisms, such as AMPAR internalization and dendritic spine shrinkage (15). Despite this, the postictal downward effects on synaptic plasticity are very robust and may act as an additional control mechanism that reduces the likelihood of immediate subsequent seizures. The downward synaptic plasticity reported here may complement other mechanisms that preclude immediate subsequent seizure, such as feedback inhibition and GABAergic signaling. In future studies, measurements of neuronal activity (calcium imaging, c-Fos, firing rate) will help to extricate the roles of homeostatic and Hebbian plasticity in the postictal period.

Along with regulating functional plasticity, AMPAR levels can modify structural plasticity, including dendritic spine density and size (9,10). Reduction in dendritic spine density has been demonstrated following stimulation protocols that induce LTD (16), chronic stimulation protocols that trigger synaptic downscaling (17), as well as neurological conditions that feature excessive neural activity like epilepsy (18). In our experiment, we did not identify changes in the overall density of dendritic spines, however, a reduction in mushroom-shaped spines was observed. As the size of the spine head is proportional to the number of postsynaptic receptors and the area of the postsynaptic density, shrinkage of mushroom spines is expected to cause weakening of synaptic transmission and to be correlated with AMPAR endocytosis (9,10). Our results support that functional, structural and molecular changes are occurring in parallel.

In sum, our data revealed that the postictal period is characterized by an LTD-like phenomenon, in which synaptic weakening, AMPARs endocytosis and shrinkage of dendritic spines are running alongside. Thus, this study provides new insights into the postictal effects of seizures on synaptic plasticity, contributing to our understanding of the neural underpinnings of the postictal seizure-suppressive state. These findings may guide the development of therapeutic strategies not only to mitigate cognitive impairments related to learning and memory, but also to reduce seizure susceptibility.

## Acknowledgements

This work was supported by São Paulo Research Foundation (FAPESP), Brazil [2018/18014-1, 2016/17882-4]. We thank Renato Meirelles e Silva, Renata Caldo Scandiuzzi and Daniela Ribeiro for technical support. We also thank Savannah May Maw for her comments on the manuscript.

## Author’s contribution

APC and JPL conceptualized the project. RNR helped APC establish the electrophysiological protocols. APC conducted the experiments and wrote the paper. APC analyzed dendritic spines and immunoblotting data. FFF helped APC with dendritic spines quantification. LRZ, DBM, RNR and APC analyzed the electrophysiological data. JPL supervised the project. All authors read, edited and approved the manuscript.

## References

1. Pottkämper JCM, Hofmeijer J, van Waarde JA, van Putten MJAM. The postictal state - What do we know?. Epilepsia. 2020;61(6):1045–1061. doi:10.1111/epi.16519

2. Kandratavicius L, Balista PA, Lopes-Aguiar C, Ruggiero RN, Umeoka EH, Garcia-Cairasco N, Bueno-Junior LS, Leite JP. Animal models of epilepsy: use and limitations. Neuropsychiatr Dis Treat. 2014 Sep 9;10:1693–705. doi: 10.2147/NDT.S50371.

3. Bragin A, Penttonen M, Buzsáki G. Termination of epileptic afterdischarge in the hippocampus. J Neurosci. 1997 Apr 1;17(7):2567–79. doi: 10.1523/JNEUROSCI.17-07-02567.1997.

4. Barr, D. S., Hoyt, K. L., Moore, S. D., & Wilson, W. A. (1997). Post-ictal depression transiently inhibits induction of LTP in area CA1 of the rat hippocampal slice. Epilepsy research, 27(2), 111–118. doi: 10.1016/s0920-1211(97)01027-9

5. Buckmaster PS, Wong EH. Evoked responses of the dentate gyrus during seizures in developing gerbils with inherited epilepsy. J Neurophysiol. 2002 Aug;88(2):783–93. doi: 10.1152/jn.2002.88.2.783.

6. Neligan A, Adan G, Nevitt SJ, Pullen A, Sander JW, Bonnett L, Marson AG. Prognosis of adults and children following a first unprovoked seizure. Cochrane Database Syst Rev. 2023 Jan 23;1(1):CD013847. doi: 10.1002/14651858.CD013847.pub2.

7. Zelig D, Goldberg I, Shor O, Ben Dor S, Yaniv-Rosenfeld A, Milikovsky DZ, Ofer J, Imtiaz H, Friedman A, Benninger F. Paroxysmal slow wave events predict epilepsy following a first seizure. Epilepsia. 2022 Jan;63(1):190–198. doi: 10.1111/epi.17110.

8. Diniz CRAF, Crestani AP. The times they are a-changin’: a proposal on how brain flexibility goes beyond the obvious to include the concepts of “upward” and “downward” to neuroplasticity. Mol Psychiatry. 2023 Mar;28(3):977–992. doi: 10.1038/s41380-022-01931-x.

9. Collingridge GL, Isaac JT, Wang YT. Receptor trafficking and synaptic plasticity. Nat Rev Neurosci. 2004 Dec;5(12):952–62. doi: 10.1038/nrn1556..

10. Kasai H, Matsuzaki M, Noguchi J, Yasumatsu N, Nakahara H. Structure-stability-function relationships of dendritic spines. Trends Neurosci. 2003 Jul;26(7):360–8. doi: 10.1016/S0166-2236(03)00162-0.

11. Citri A, Malenka RC. Synaptic plasticity: multiple forms, functions, and mechanisms. Neuropsychopharmacology. 2008 Jan;33(1):18–41. doi: 10.1038/sj.npp.1301559.

12. Hesse GW, Teyler TJ. Reversible loss of hippocampal long term potentiation following electronconvulsive seizures. Nature. 1976 Dec 9;264(5586):562–4. doi: 10.1038/264562a0.

13. Liu X, Muller RU, Huang LT, Kubie JL, Rotenberg A, Rivard B, Cilio MR, Holmes GL. Seizure-induced changes in place cell physiology: relationship to spatial memory. J Neurosci. 2003 Dec 17;23(37):11505–15. doi: 10.1523/JNEUROSCI.23-37-11505.2003.

14. Turrigiano G. Homeostatic synaptic plasticity: local and global mechanisms for stabilizing neuronal function. Cold Spring Harb Perspect Biol. 2012 Jan 1;4(1):a005736. doi: 10.1101/cshperspect.a005736..

15. Turrigiano GG. The dialectic of Hebb and homeostasis. Philos Trans R Soc Lond B Biol Sci. 2017 Mar 5;372(1715):20160258. doi: 10.1098/rstb.2016.0258.

16. Zhou Q, Homma KJ, Poo MM. Shrinkage of dendritic spines associated with long-term depression of hippocampal synapses. Neuron. 2004 Dec 2;44(5):749–57. doi: 10.1016/j.neuron.2004.11.011.

17. Goold CP, Nicoll RA. Single-cell optogenetic excitation drives homeostatic synaptic depression. Neuron. 2010 Nov 4;68(3):512–28. doi: 10.1016/j.neuron.2010.09.020.

18. Swann JW, Al-Noori S, Jiang M, Lee CL. Spine loss and other dendritic abnormalities in epilepsy. Hippocampus. 2000;10(5):617–25. doi: 10.1002/1098-1063(2000)10:5<617::AID-HIPO13>3.0.CO;2-R.

